# ALS-causing SOD1 mutations H46R and G85R form similar novel amyloid fibril structures and promote ferroptosis in cells

**DOI:** 10.1101/2023.06.13.544727

**Authors:** Li-Qiang Wang, Yeyang Ma, Mu-Ya Zhang, Han-Ye Yuan, Xiang-Ning Li, Wencheng Xia, Kun Zhao, Xi Huang, Jie Chen, Liangyu Zou, Dan Li, Zhengzhi Wang, Weidong Le, Cong Liu, Yi Liang

## Abstract

More than two hundred genetic mutations of Cu, Zn-superoxide dismutase (SOD1) have been identified in amyotrophic lateral sclerosis (ALS), a neurodegenerative disease characterized by the selective death of motor neurons through ferroptosis. Two ALS-causing SOD1 mutations, H46R and G85R, are metal-binding region mutants with reduced affinity for metal ions. Here, we generated amyloid fibrils from the apo forms of H46R and G85R under reducing conditions and determined their structures using cryo-EM. We built models for the fibril cores, comprising residues 85−153 for H46R and 82−153 for G85R. These mutations disrupt crucial interactions in the wild-type SOD1 fibril, resulting in amyloid fibrils with distinct structures compared to the wild-type fibril. Remarkably, H46R and G85R form similar novel amyloid fibril structures. The fibril cores display a serpentine fold containing seven or eight β-strands, which are stabilized by a hydrophobic cavity. In the G85R fibril core, Arg85 and Asp101 form a salt bridge for stabilization. We demonstrate that fibril seeds from H46R and G85R cause more severe mitochondrial impairment and significantly promote ferroptosis in neuronal cells, compared with those from wild-type SOD1. Our findings reveal how different SOD1 mutations can result in similar amyloid fibril structures and contribute to ALS pathology.

## Introduction

Amyotrophic lateral sclerosis (ALS), also known as motor neuron disease, is a progressive, devastating neurodegenerative disease characterized by selective death of motor neurons in the brain and spinal cord^1–6^ through ferroptosis^7–9^. Ferroptosis, an iron-dependent form of nonapoptotic cell death^7, 9–11^, is regulated by glutathione peroxidase 4 (GPX4)^12, 13^, and GPX4 is downregulated in the spinal cords of patients with ALS^7, 8^. However, the molecular mechanism underlying the mediation of ferroptosis in ALS remains unclear. Approximately 90% of ALS cases are sporadic while about 10% are familial and are typically passed down with inheritance^1–4, 14–16^. There are now more than 50 genes that have been implicated in ALS pathogenesis^1^. Among them, the *sod1* gene, the first gene to be associated with a familial form of ALS^17^, is the second most common cause of the disease^1, 15, 16^. About 2−6% of cases of ALS are caused by mutations in the antioxidant enzyme Cu, Zn-superoxide dismutase (SOD1)^1, 2, 4–6, 14–16^. Strikingly, more than two hundred genetic mutations of SOD1 have been identified in the familial form of ALS^1–5, 14, 17–37^ (https://alsod.iop.kcl.ac.uk/). These mutations have remarkably diverse effects on the structure, activity, and stability of the native state of SOD1 (refs. ^1–3, 14, 20, 27, 29, 30, 33, 36, 37^). The misfolding and aggregation of these mutations in motor neuron cells play a crucial role in the etiology and pathogenesis of ALS^1, 2–4, 14, 15, 20–24, 27, 28, 30, 32, 33^. Motor neuronal degeneration in ALS has been implicated to be caused by toxic gain of function of mutant SOD1 in addition to loss-of-function^1, 3, 5, 6, 14, 30, 31, 38–43^. It is, however, currently unknown whether different ALS-causing SOD1 mutations produce distinct SOD1 strains that influence the evolution of the disease^1, 44, 45^ and whether they promote SOD1 aggregation by fundamentally distinct mechanisms^1–4, 22–24, 27–30, 34, 38, 44, 45^.

Since SOD1 was found to be associated with familial ALS in 1993 (ref. ^17^), great efforts have been dedicated to unravel the mysteries of the atomic structure of SOD1 aggregates^1, 46–50^ and SOD1 strains^1, 44, 45^. Recently, we reported a cryo-EM structure of the amyloid fibril formed by the disulfide-reduced, apo form of full-length wild-type human SOD1 featuring an in-register intramolecular β sheet architecture^46^, which provides structural insights into the conversion of SOD1 from its immature form into an aggregated form during ALS pathogenesis. However, despite three decades of investigation^1–5, 17–46^, the molecular mechanisms by which mutations in SOD1 cause the familial form of ALS remain a mystery.

Various SOD1 mutations include metal-binding region mutants, such as H46R, H46D, G85R, D125H, and S134N, and wild-type-like mutants, such as A4V, D90A, G93A, D101G, and D101N^1–5, 14, 17–37, 44–46^. In this study, we focus specifically on two ALS-causing SOD1 mutations, H46R and G85R, because of the following reasons. First, H46R and G85R are metal-binding region mutants with reduced affinity for metal ions (zinc in the case of G85R and copper and zinc for H46R)^2, 14, 18, 19, 29, 36^; the aggregates or inclusions formed by H46R and G85R exhibit prion-like properties^34, 35^. Second, transgenic mice expressing H46R and G85R develop a similar ALS-like phenotype comprising of paralysis and muscle loss at several months of age^1, 3, 21, 22, 31–33, 35^. However, the mechanism behind the phenomenon remains unclear. Third, the atomic structure of wild-type SOD1 fibrils has shown that several familial mutations including H46R, H46D, G85R, D101G, and D101N may disrupt crucial interactions (salt bridges) in the cytotoxic SOD1 fibril structure^46^, but whether ALS-causing SOD1 mutations H46R and G85R form similar novel amyloid fibril structures is unknown.

Here, we generated homogeneous amyloid fibrils from the apo forms of two ALS-causing SOD1 mutations, H46R and G85R, under reducing conditions and determine the atomic structure by using cryo-EM. We demonstrate that H46R and G85R form similar novel amyloid fibril structures and significantly promote ferroptosis in neuronal cells. Our findings provide structural insights into the molecular mechanisms by which mutations in SOD1 promote ferroptosis regulated by GPX4 and cause the familial form of ALS.

## Results

### Comparison of the cryo-EM structures of the H46R fibril and the G85R fibril

We first treated the apo forms of H46R and G85R with 5 mM tris (2-carboxyethyl) phosphine (TCEP). TCEP, a highly stable disulfide-reducing agent, can be used to generate a reduced state that is able to mimic physiological reducing environments^46^. We produced amyloid fibrils from recombinant, full-length apo human SOD1 (residues 1 to 153) with H46R mutation or G85R mutation overexpressed in *Escherichia coli*, by incubating the purified apoproteins in 20 mM tris-HCl buffer (pH 7.4) containing 5 mM TCEP and shaking at 37 °C for 40−48 h (see methods). Amyloid fibrils formed by H46R and G85R under such reducing conditions were concentrated to ∼30 μM in a centrifugal filter (Millipore) and examined by electron microscopy without further treatment.

Negative-staining transmission electron microscopy (TEM) images showed that the apo forms of ALS-causing SOD1 mutations H46R and G85R formed homogeneous and unbranched fibrils under reducing conditions (Extended Data Fig. 1a,b). We then compared the images of amyloid fibrils assembled from H46R and G85R by atomic force microscopy (AFM) (Extended Data Fig. 1c,d) and determined the atomic structures of the H46R fibril and the G85R fibril by cryo-EM (Table 1 and Figs. 1 and 2). The AFM images, cryo-EM micrographs, and two-dimensional (2D) class average images using RELION3.1 (ref. ^51^) showed that both the H46R fibril and the G85R fibril were composed of a single protofilament with a left-handed helical twist (Extended Data Fig. 1c−f) and arranged in a staggered manner (Extended Data Fig. 2a,b). The helical pitch was 170 ± 9 nm for the H46R fibril (Extended Data Fig. 1c) and 168 ± 7 nm for the G85R fibril (Extended Data Fig. 1d). The fibrils were morphologically homogeneous (Extended Data Fig. 1c−f), showing a fibril full width of 9.8 ± 1.0 nm for the H46R fibril (Extended Data Fig. 1c) and 10.0 ± 1.4 nm for the G85R fibril (Extended Data Fig. 1d).

**Fig. 1.**
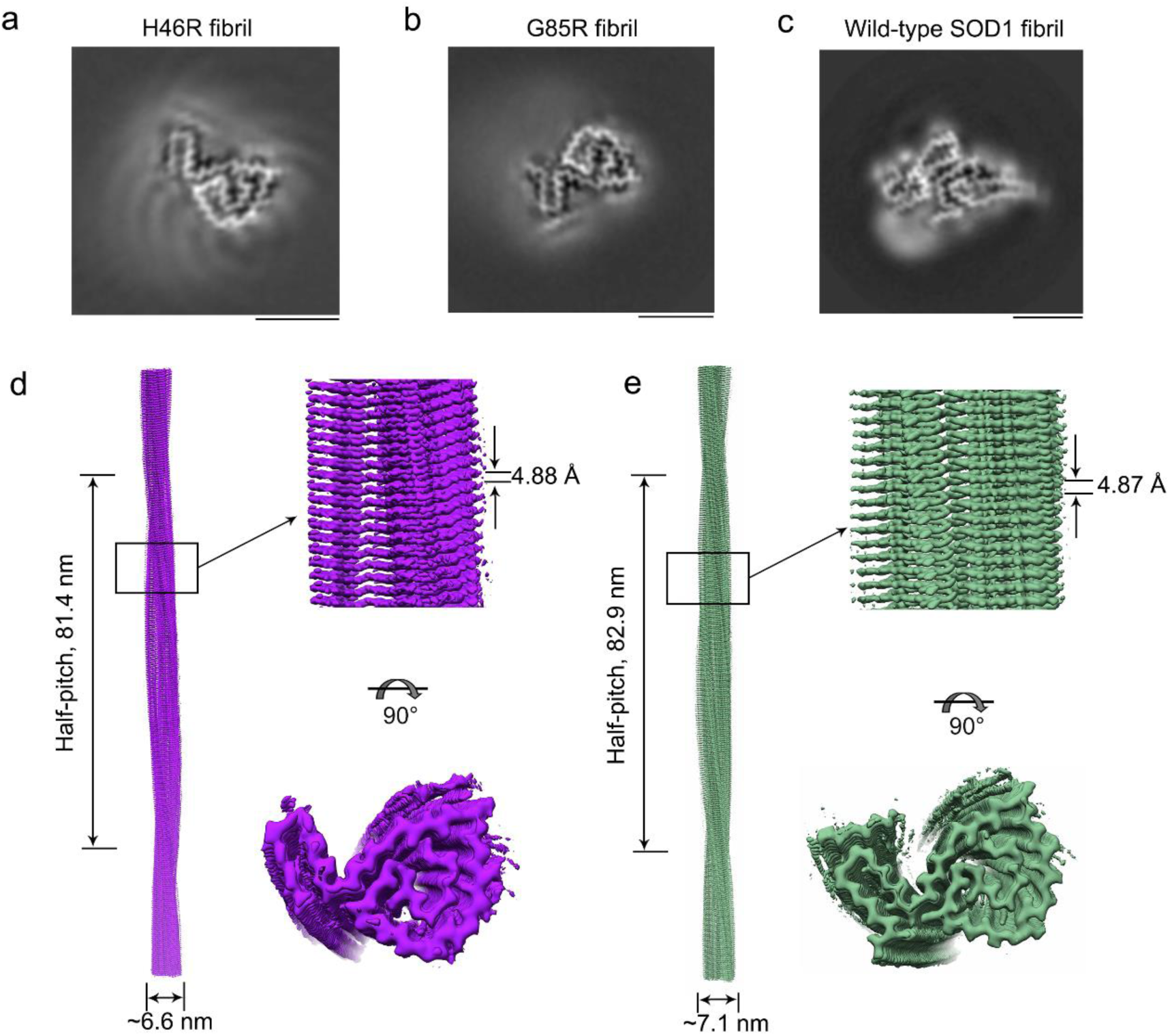
Comparison of the cryo-EM structures of the H46R fibril and the G85R fibril. **a**,**b**, Cross-sectional view of the 3D map of the H46R fibril (**a**) or the G85R fibril (**b**) showing a protofilament comprising a C-terminal segment. **c**, Cross-sectional view of the 3D map of the wild-type SOD1 fibril, however showing a protofilament comprising not only a C-terminal segment (right) but also a N-terminal segment (left) with an unstructured flexible region (bottom)^50^. Scale bars, 5 nm. **d**,**e**, 3D map of the H46R fibril (**d**) or the G85R fibril (**e**) showing a single protofilament (in purple for **d** and green for **e**) intertwined into a left-handed helix, with a fibril core width of ∼6.6 (**d**) or ∼7.1 (**e**) nm and a half-helical pitch of 81.4 (**d**) or 82.9 (**e**) nm (left). Enlarged section of the H46R fibril (**d**) or the G85R fibril (**e**) showing a side view of the density map (top right). Close-up view of the density map in the left showing that the subunit in a protofilament stacks along the fibril axis with a helical rise of 4.88 (**d**) or 4.87 (**e**) Å (top right). Top view of the density map of the H46R fibril (**d**) or the G85R fibril (**e**) (bottom right).

**Fig. 2.**
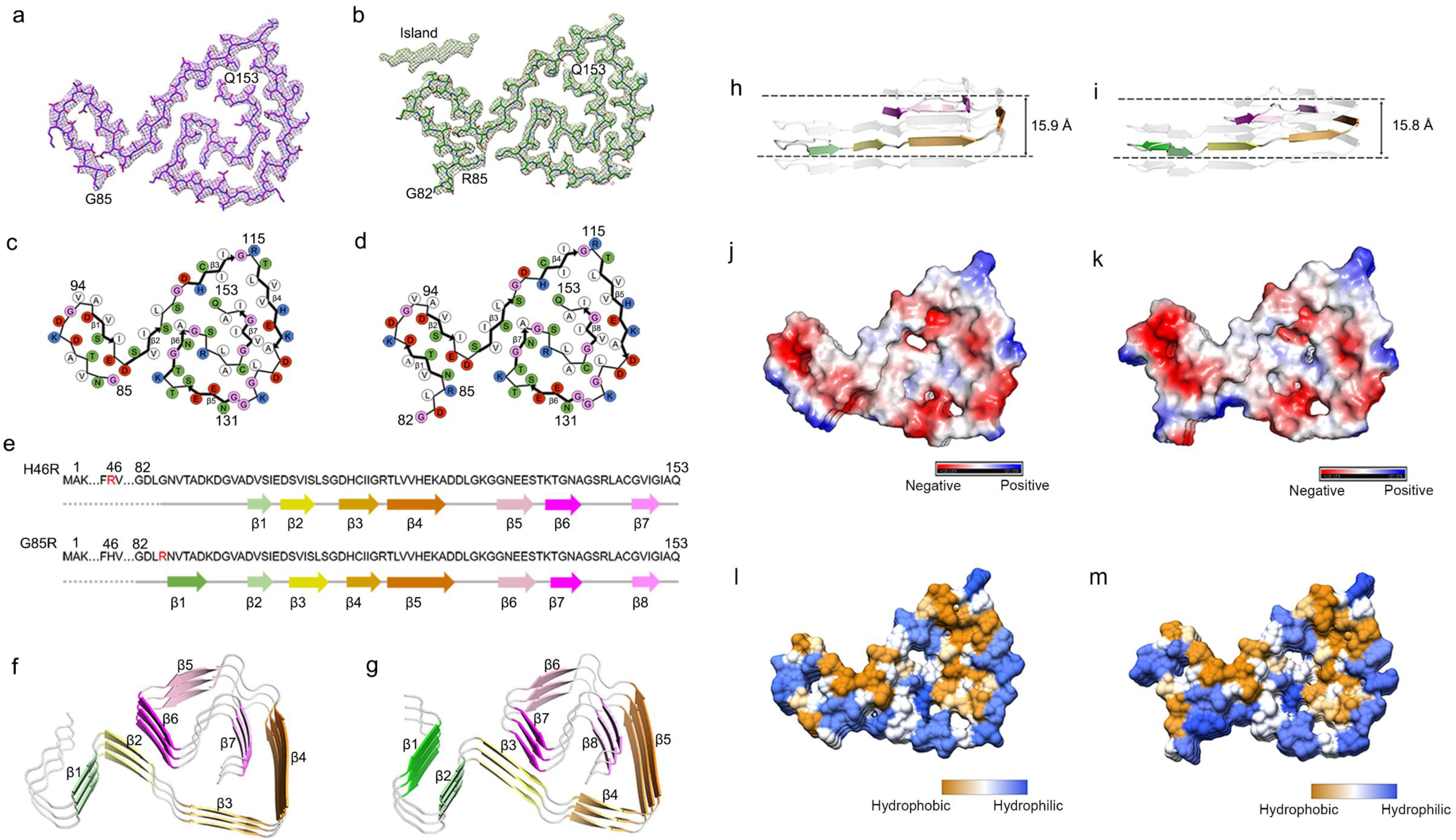
ALS-causing SOD1 mutations H46R and G85R form similar novel amyloid fibril structures. **a**, Cryo-EM map of the H46R fibril with the atomic model overlaid. The H46R fibril core comprises a C-terminal segment (residues 85 to 153) colored purple. **b**, Cryo-EM map of the G85R fibril with the atomic model overlaid. The G85R fibril core comprises a C-terminal segment (residues 82 to 153) colored green. One density (termed an island) flanking the protofilament, which is also colored green. **c**,**d**, Schematic view of the H46R fibril core (**c**) or the G85R fibril core (**d**). Residues are colored as follows: white, hydrophobic; green, polar; red and blue, negatively charged and positively charged, respectively; and magenta, glycine. β strands are indicated with bold lines. **e**, Sequence of the H46R fibril core comprising residues 85−153 from the full-length human H46R SOD1 (1 to 153) with the observed seven β strands colored light green (β1), yellow (β2), gold (β3), orange (β4), pink (β5), magenta (β6), and light magenta (β7) in the C-terminal segment (top). Dotted line corresponds to residues 1−84 not modeled in the cryo-EM density. Sequence of the G85R fibril core comprising residues 82−153 from the full-length human G85R SOD1 (1 to 153) with the observed eight β strands colored green (β1), light green (β2), yellow (β3), gold (β4), orange (β5), pink (β6), magenta (β7), and light magenta (β8) in the C-terminal segment (bottom). Dotted line corresponds to residues 1−81 not modeled in the cryo-EM density. ALS-causing mutation sites, Arg46 and Arg85, are highlighted in red. **f**,**g**, Ribbon representation of the structure of an H46R fibril core (**f**) or an G85R fibril core (**g**) containing three molecular layers and a C-terminal segment. **h**,**i**, As in **f**,**g**, but viewed perpendicular to the helical axis, revealing that the height of one layer along the helical axis is 15.9 Å or 15.8 Å. **j**,**k**, Electrostatic surface representation of the structure of an H46R fibril core (**j**) or an G85R fibril core (**k**) containing three molecular layers and a C-terminal segment. **l**,**m**, Hydrophobic surface representation of the structure of an H46R fibril core as in **f** or an G85R fibril core as in **g**. The surface of the H46R fibril core or the G85R fibril core is shown according to the electrostatic properties (red, negatively charged; blue, positively charged) (**j** and **k**) or the hydrophobicity (yellow, hydrophobic; blue, hydrophilic) (**l** and **m**) of the residues.

**Table 1.** Cryo-EM data collection, refinement, and validation statistics.

|  | **H46R fibril**<br>**(EMD-35460, PDB**<br>**8IHV)** | **G85R fibril**<br>**(EMD-35459, PDB**<br>**8IHU)** |
| --- | --- | --- |
| **Data collection and processing** |  |  |
| Magnification | 105,000 | 105,000 |
| Voltage (kV) | 300 | 300 |
| Camera | Gatan K3<br>(Krios G4) | Gatan K3<br>(Krios G4) |
| Frame exposure time (s) | 0.08 | 0.08 |
| Movie frames (*n*) | 40 | 40 |
| Electron exposure (e<sup>-</sup>/Å<sup>2</sup>) | 60 | 60 |
| Defocus range (μm) | −2.0 to −1.2 | −2.0 to −1.2 |
| Pixel size (Å) | 0.84 | 0.84 |
| Symmetry imposed | C1 | C1 |
| Box size (pixel) | 400 | 400 |
| Inter-box distance (Å) | 33.6 | 33.6 |
| Micrographs collected (*n*) | 8,680 | 6,366 |
| Segments extracted (*n*) | 1,062,069 | 626,757 |
| Segments after Class2D (*n*) | 235,369 | 305,672 |
| Segments after Class3D (*n*) | 59,857 | 53,707 |
| Map resolution (Å) | 3.11 | 2.97 |
| FSC threshold | 0.143 | 0.143 |
| Map resolution range (Å) | 3.01–4.83 | 2.87–4.79 |
| **Refinement** |  |  |
| Initial model used | De novo | De novo |
| Model resolution (Å) | 3.11 | 2.97 |
| FSC threshold | 0.143 | 0.143 |
| Model resolution range (Å) | 3.11 | 2.97 |
| Map sharpening *B* factor (Å<sup>2</sup>) | −78.38 | −82.56 |
| Model composition |  |  |
| Nonhydrogen atoms | 1,446 | 1,527 |
| Protein residues | 207 | 216 |
| Ligands | 0 | 0 |
| *B* factors (Å<sup>2</sup>) |  |  |
| Protein | 81.95 | 52.90 |
| R.m.s. deviations |  |  |
| Bond lengths (Å) | 0.005 | 0.005 |
| Bond angles (°) | 0.722 | 0.713 |
| **Validation** |  |  |
| MolProbity score | 2.39 | 2.25 |
| Clashscore | 12.57 | 9.90 |
| Poor rotamers (%) | 0 | 0 |
| Ramachandran plot |  |  |
| Favored (%) | 76.12 | 80.00 |
| Allowed (%) | 23.88 | 20.00 |
| Disallowed (%) | 0 | 0 |

Using helical reconstruction in RELION3.1 (ref. ^51^), we determined density maps of the ordered cores of H46R fibril and G85R fibril, with overall resolutions of 3.11 Å and 2.97 Å, respectively, which featured well-resolved side-chain densities and clearly separated β strands along the fibril axis (Fig. 1a,b and Extended Data Fig. 3a,b). Cross-sectional views of the 3D maps of the H46R fibril and the G85R fibril showed a protofilament comprising a C-terminal segment (Fig. 1a,b). Cross-sectional view of the 3D map of the wild-type SOD1 fibril, however, showed a protofilament comprising not only a C-terminal segment but also a N-terminal segment, with an unstructured flexible region in between (Fig. 1c). 3D maps of the H46R fibril and the G85R fibril showed a single protofilament intertwined into a left-handed helix, with fibril core widths of ∼6.6 nm and ∼7.1 nm and half-helical pitches of 81.4 nm or 82.9 nm, respectively (Fig. 1d,e). The subunits within the protofilaments of H46R and G85R stacked along the fibril axis with helical rises of 4.88 Å and 4.87 Å and twists of −1.079° and −1.058°, respectively (Fig. 1d,e). Together, the data showed that under reducing conditions, H46R fibril and G85R fibril displayed similar novel structures.

**Fig. 3.**
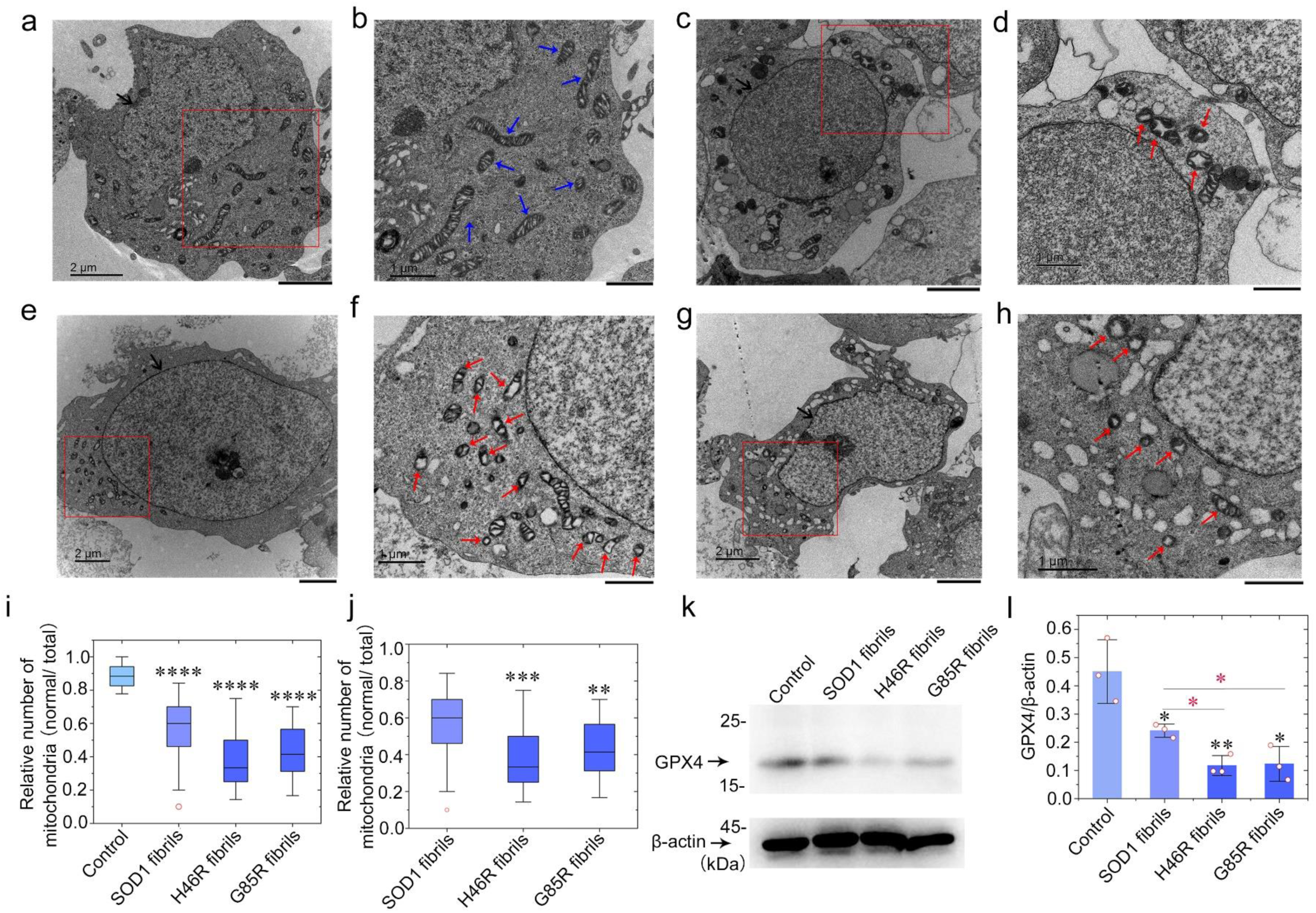
Fibril seeds from H46R and G85R cause much more severe mitochondrial impairment and significantly promote ferroptosis in neuronal cells, compared with those from wild-type SOD1. **a**−**h**, SH-SY5Y neuroblastoma cells were cultured for 1 day and then incubated with 0 μM SOD1 fibril seeds (**a** and **b**), 10 μM wild-type SOD1 fibril seeds (**c** and **d**), 10 μM H46R fibril seeds (**e** and **f**), and 10 μM G85R fibril seeds (**g** and **h**), respectively, for 3 days. The enlarged regions (**b**, **d**, **f**, and **h**) show 4-fold, 5-fold, 10-fold, and 9-fold enlarged images from **a**, **c**, **e**, and **g**, respectively, and display the detailed structures of mitochondria in cells. Nuclei are highlighted using black arrows (**a**, **c**, **e**, and **g**). **b**, Normal mitochondria in SH-SY5Y cells are highlighted by blue arrows. A number of mitochondria in the cells (**d**) and most of mitochondria in the cells (**f** and **h**) partly became smaller than normal mitochondria with increased mitochondrial membrane density and reduction of mitochondria crista, indicating ferroptosis, and partly became swollen and vacuolized, which is highlighted by red arrows. Samples were negatively stained using 2% uranyl acetate and lead citrate. The scale bars represent 2 μm (**a**, **c**, **e**, and **g**) and 1 μm (**b**, **d**, **f**, and **h**), respectively. **i**,**j**, Box plots showing the quantification of TEM images in *n* = 30 SH-SY5Y cells examined over 3 independent experiments. The boxes (blue) extend from the 25th to 75th percentile (quantiles 1 and 3). Minima, maxima, centre, and bounds of box represent quantile 1 minus 1.5 × interquartile range, quantile 3 plus 1.5 × interquartile range, median (black line), and quantiles 1 and 3, respectively. Bounds of whiskers (minima and maxima) and outliers (open red circles). Cells treated with 20 mM Tris-HCl buffer (pH 7.4) containing 5 mM TCEP for 3 days (**i**) and cells incubated with 10 μM wild-type SOD1 fibril seeds (**j**) were used as controls, respectively. **k**, SH-SY5Y cells were cultured for 1 day and then incubated with 0 μM SOD1 fibril seeds, 10 μM wild-type SOD1 fibril seeds, 10 μM H46R fibril seeds, and 10 μM G85R fibril seeds, respectively, for 3 days. The cell lysates from the above cells were probed by the anti-GPX4 antibody and anti-β-actin antibody, respectively. **l**, The relative amount of GPX4 in the above cell lines (open red circles shown in scatter plots) was determined as a ratio of the density of GPX4 band over the density of β-actin band in cell lysates and expressed as the mean ± S.D. (with error bars) of values obtained in three independent experiments. *p*-values were determined using two-sided Student’s *t*-test. Values of *p* < 0.05 indicate statistically significant differences. The following notation is used throughout: \**p* < 0.05; \*\**p* < 0.01; \*\*\**p* < 0.001; and \*\*\*\**p* < 0.0001 relative to control. Source data are provided as a Source Data file.

### ALS-causing SOD1 mutations H46R and G85R form similar novel amyloid fibril structures

We unambiguously built a structure model of H46R fibril comprising a C-terminal segment (residues 85 to 153) at 3.11 Å and that of G85R fibril comprising a C-terminal segment (residues 82 to 153) at 2.97 Å (Fig. 2). Side chain densities for many residues in the H46R fibril and most residues in the G85R fibril had high local resolution (3.00−3.125 Å) (Extended Data Fig. 3c,d). Side chains for the residues in the fibril cores of H46R and G85R can be well accommodated into the density maps (Fig. 2a,b). We observed one unidentified flanking the protofilament in the G85R fibril, termed an island (Fig. 2b). This island is located on the opposing side of hydrophobic side chains of Val94 and Ala95 in the G85R fibril (Fig. 2b), and is reminiscent of the island observed in the structure of an amyloid fibril formed by full-length human prion protein with E196K mutation, a genetic Creutzfeldt-Jakob disease–related mutation^52^. The exteriors of the fibril cores of H46R and G85R are partly hydrophilic, carrying many negatively charged or positively charged residues, whereas side chains of most hydrophobic residues are mainly located in the interiors of the H46R/G85R fibril fold (Fig. 2c−m). A hydrophobic core (Fig. 2c,d,l,m), three hydrogen bonds (Extended Data Figs. 4a,b and 5a−f), a salt bridge (Extended Data Fig. 4c,d), and a very compact fold (Fig. 2c,d,f,g) help stabilize the fibril cores, as described in detail below.

**Fig. 4.**
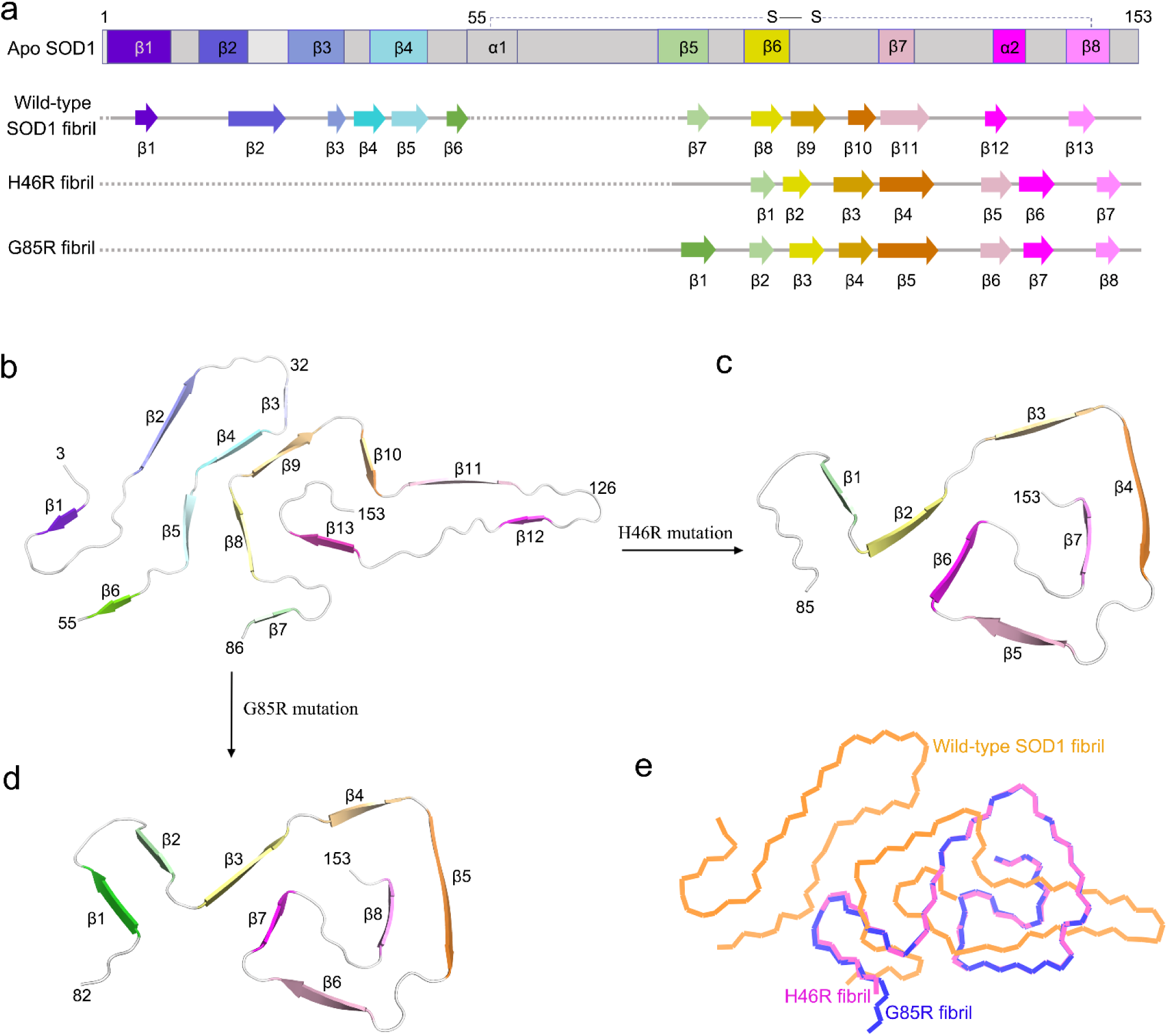
Comparison of the structures of the apo form of SOD1, the wild-type SOD1 fibril, the H46R fibril, and the G85R fibril. **a**, Sequence alignment of the full-length apo human SOD1 (1 to 153) monomer (PDB 1HL4)^62^ with eight β-strands colored violet (β1), blue (β2), light blue (β3), light cyan (β4), light green (β5), yellow (β6), pink (β7), and light magenta (β8), two α-helices colored gray (α1) and magenta (α2), and a single disulfide bond between Cys57 in α1 and Cys146 in β8. Sequence alignment of the wild-type SOD1 fibril core comprising residues 3−55 and 86−153 from the full-length wild-type human SOD1 (PDB 7VZF)^50^ with the observed thirteen β strands colored violet (β1), blue (β2), light blue (β3), cyan (β4), light cyan (β5), green (β6), light green (β7), yellow (β8), gold (β9), orange (β10), pink (β11), magenta (β12), and light magenta (β13). Dotted lines correspond to residues 1−2 and residues 56−85, which were not modeled in the cryo-EM density. Sequence alignment of the H46R fibril core comprising residues 85−153 from the full-length human H46R SOD1 (1 to 153) with the observed seven β strands colored light green (β1), yellow (β2), gold (β3), orange (β4), pink (β5), magenta (β6), and light magenta (β7) in the C-terminal segment. Dotted line corresponds to residues 1−84 not modeled in the cryo-EM density. Sequence alignment of the G85R fibril core comprising residues 82−153 from the full-length human G85R SOD1 (1 to 153) with the observed eight β strands colored green (β1), light green (β2), yellow (β3), gold (β4), orange (β5), pink (β6), magenta (β7), and light magenta (β8) in the C-terminal segment (bottom). Dotted line corresponds to residues 1−81 not modeled in the cryo-EM density. **b**−**d**, Ribbon representation of the structures of a wild-type SOD1 fibril core (PDB 7VZF)^50^ (**b**), an H46R fibril core (**c**), and an G85R fibril core (**d**), all of which contain one molecular layer and a monomer. **e**, Overlay of the structures of a wild-type SOD1 fibril core (orange) (PDB 7VZF)^50^, an H46R fibril core (magenta), and an G85R fibril core (blue).

Hydrophobic side chains of Ile112, Leu117, Val119, Ala123, Leu126, Leu144, Val148, Ile149, Ile151, and Ala152 are found to locate in the interior of H46R fibrils (Fig. 2c) or G85R fibrils (Fig. 2d) to form a stable hydrophobic core. In sharp contrast, hydrophilic side chains of Thr88, Asp90, Asp96, Ser98, Glu100, and Asp101 are found to locate in the interior of H46R fibrils to form a hydrophilic cavity (Fig. 2c), and hydrophilic side chains of Arg85, Asn86, Thr88, Asp90, Asp96, Ser98, Glu100, and Asp101 are found to locate in the interior of G85R fibrils to form a hydrophilic cavity (Fig. 2d). Hydrophilic side chains of Ser105, Ser107, His110, Ser142, and Gln153 are found to locate in the interior of H46R fibrils (Fig. 2c) or G85R fibrils (Fig. 2d) to form the second hydrophilic cavity. Hydrophilic side chains of Glu132, Ser134, Thr137, Asn139, and Arg143 are found to locate in the interior of H46R fibrils (Fig. 2c) or G85R fibrils (Fig. 2d) to form the third hydrophilic cavity.

Importantly, Arg85 and Asp101 form a salt bridge with a distance of 2.5 Å to stabilize the G85R fibril core (Extended Data Fig. 4c,d), whereas Asn86 and Asp101 form a hydrogen bond with a distance of 2.6 Å to stabilize the H46R fibril core (Extended Data Fig. 4a,b). Two pairs of amino acids (His110 and Gln153; and Ser134 and Arg143) form two or three hydrogen bonds to stabilize the fibril cores (Extended Data Fig. 5a−f). The fibril core structures of H46R and G85R only comprise a C-terminal segment containing residues 85−153 and 82−153, respectively (Fig. 2a−g). The wild-type SOD1 fibril core structure, however, comprises not only a C-terminal segment (residues 3 to 55) but also a N-terminal segment (residues 86 to 153), with an unstructured flexible region in between^46^. Thus, the H46R mutation and the G85R mutation disrupt crucial salt bridges in the wild-type SOD1 fibril and make important interactions in the fibril cores (Extended Data Figs. 4 and 5), resulting in amyloid fibrils with distinct structures compared to the wild-type fibril (Figs. 1 and 2).

The fibril cores feature a very compact fold and exhibit a serpentine fold containing seven or eight β-strands stabilized by a hydrophobic cavity (Fig. 2c,d,f,g,l,m). Seven β-strands (β1 to β7) and eight β-strands (β1 to β8) are present in the fibril core structures of H46R and G85R, respectively (Fig. 2c−g). The height of one layer of the H46R fibril core (or the G85R fibril core) along the helical axis is 15.9 Å (or 15.8 Å), which is the distance between the highest point in the loop between β6 and β7 (or between β7 and β8) and the lowest point in the loop between β1 and β2 (Fig. 2h,i). Together, these results demonstrate that H46R and G85R form similar novel amyloid fibril structures.

### Fibril seeds from H46R and G85R significantly promote ferroptosis in neuronal cells

SOD1 fibrils produced under reducing conditions have functions to induce mitochondria damage^46, 53^. Ferroptosis has functional importance in mediating motor neuron death in ALS^7–9^. Specifically, mitochondria in ferroptotic cells appear smaller than normal and show increased mitochondrial membrane density, decreased mitochondrial crista, and rupture of the outer membrane^10, 54, 55^. Given that two ALS-causing SOD1 mutations, H46R and G85R, do form similar amyloid fibril structures (Fig. 2), we predicted that these two mutations might perform similar functions implicated in ALS, exhibiting similar ability to induce mitochondria damage and activate ferroptosis in neuronal cells.

We next used ultrathin section TEM and western blotting to test this hypothesis. The morphology of normal mitochondria in SH-SY5Y neuroblastoma cells incubated with 0 μM SOD1 fibril seeds, which are highlighted by blue arrows, was tubular or round (Fig. 3a,b). 10 μM wild-type SOD1 fibril seed treatment caused serious mitochondrial impairment and induced ferroptosis in SH-SY5Y cells (Fig. 3c,d). Importantly, 10 μM H46R fibril seed treatment (Fig. 3e,f) and 10 μM G85R fibril seed treatment (Fig. 3g,h) caused severe mitochondrial impairment and promoted ferroptosis in SH-SY5Y cells. A number of mitochondria in the cells (∼40%) (Fig. 3d,i,j, for wild-type SOD1 fibril seed-treated cells) and most of mitochondria in the cells (∼70%) (Fig. 3f,h,i,j, for H46R fibril seed- and G85R fibril seed-treated cells) partly became smaller than normal mitochondria with increased mitochondrial membrane density and reduction of mitochondria crista, indicating ferroptosis, and partly became swollen and vacuolized, which is highlighted by red arrows. A significantly lower number of normal mitochondria was observed in SH-SY5Y cells treated by wild-type SOD1 fibril seeds, H46R fibril seeds, and G85R fibril seeds than did in control cells treated by Tris-HCl buffer containing TCEP (*p* = 3.3 × 10^-12^, 5.6 × 10^-21^, and 5.9 × 10^-22^, respectively) (Fig. 3i). Importantly, a significantly lower number of normal mitochondria was observed in SH-SY5Y cells treated by H46R fibril seeds and G85R fibril seeds than did in control cells treated by wild-type SOD1 fibril seeds (*p* = 0.00020 and 0.0016, respectively) (Fig. 3j). To gain a quantitative understanding of how SOD1 mutations regulate neuronal cell ferroptosis, we detected a master regulator of ferroptosis, GPX4 (ref. ^7,12, 13^), in the above cells using the anti-GPX4 antibody (Fig. 3k). Upon incubation with 10 μM fibril seeds for 3 days, GPX4 was downregulated in SH-SY5Y cells treated by wild-type SOD1 fibril seeds where GPX4 protein levels were reduced by ∼46% (*p* = 0.0346) (Fig. 3k,l). Intriguingly, we observed that GPX4 was strongly downregulated in SH-SY5Y cells treated by H46R fibril seeds and G85R fibril seeds for 3 days, and GPX4 protein levels were reduced by ∼74% in H46R fibril seed-treated cells (*p* = 0.0081) and ∼73% in G85R fibril seed-treated cells (*p* = 0.0116), compared to control cells treated by Tris-HCl buffer containing TCEP when normalized to β-actin (Fig. 3k,l). Strikingly, GPX4 protein levels in H46R fibril seed- and G85R fibril seed-treated cells were significantly decreased compared with control cells treated by wild-type SOD1 fibril seeds (*p* = 0.0072 and 0.0367, respectively), indicating that treatment of cells with 10 μM H46R fibril seeds or 10 μM G85R fibril seeds significantly promoted ferroptosis in SH-SY5Y cells (Fig. 3l). These results demonstrate that the fibril seeds from H46R and G85R cause more severe mitochondrial impairment and significantly promote ferroptosis in neuronal cells in a similar way, compared with those from wild-type SOD1.

Altogether these data demonstrate that ALS-causing SOD1 mutations H46R and G85R form similar amyloid fibril structures and strongly suggest that these different SOD1 mutations exhibit similar ability to induce mitochondria damage and activate ferroptosis in neuronal cells, contributing to ALS pathology.

## Discussion

Mutations in SOD1 account for about 2−6% of all ALS^1, 2, 14–16, 56^. Because familial mutations in SOD1, such as H46R and G85R, are involved in the pathogenesis of the motor neuron disease ALS where it is observed to form intracellular fibrillar inclusions^3, 14, 20, 32, 33, 35, 57^, it has generally been thought that these proteinaceous inclusions could be responsible for neuronal cell death in patients with ALS^3, 14, 35^. Here, we compared the structures of apo SOD1, the wild-type SOD1 fibril, the H46R fibril, and the G85R fibril (Fig. 4). Notably, the SOD1 molecule adopts largely distinctive secondary and tertiary structures in three different SOD1 structures (apo SOD1, the wild-type SOD1 fibril, and the H46R/G85R fibril) (Fig. 4a−d), highlighting the phenotypic diversity of SOD1 in physiological and fibrillar states. The full-length apo human SOD1 monomer contains eight β-strands (β1 to β8), two α-helices (α1 and α2), and a single disulfide bond between Cys57 in α1 and Cys146 in β8 (ref. ^58^) (Fig. 4a). Once it folds into its fibrillar form under reducing conditions, the SOD1 subunit undergoes a totally conformational rearrangement. The wild-type human SOD1 fibril core contains six β-strands (β1 to β6) by its N-terminal segment (residues 3 to 55) and seven β-strands (β7 to β13) by its C-terminal segment (residues 86 to 153), exhibiting an in-register intramolecular β sheet architecture^46^ (Fig. 4a,b). In sharp contrast, the fibril cores of H46R and G85R only comprise a C-terminal segment with residues 85−153 and 82−153 containing seven β-strands (β1 to β7) and eight β-strands (β1 to β8), respectively (Fig. 4a,c,d). The H46R mutation and the G85R mutation disrupt crucial interactions in SOD1 fibrils and results in a rearrangement of the overall structure, resulting in amyloid fibrils with distinct structures compared to the wild-type fibril (Fig. 4b−e). H46R and G85R form similar amyloid fibril structures with an r.m.s.d of 0.273 Å (55 to 55 Cα atoms), while the wild-type SOD1 fibril could hardly align with the H46R fibril and the G85R fibril with r.m.s.d of 16.775 Å (68 to 68 Cα atoms) and 16.776 Å (68 to 68 Cα atoms), respectively (Fig. 4e).

Remarkably, ALS-causing SOD1 mutations H46R and G85R form similar amyloid fibril structures (Fig. 4c,d) and exhibit similar ability to induce mitochondria damage and activate ferroptosis in neuronal cells (Fig. 3), highlighting the degeneracy properties of these familial mutations in an amyloid state. Very recently, Wilkinson and co-workers have suggested that amyloid fibrils formed by three disease-causing mutations in β_2_-microglobulin (ΔN6, D76N, and V27M) at more physiological pH 6.2 share a common amyloid fold but with different arrangements of these folds within their fibril architectures^59^. This is partly compatible with our model, wherein different ALS-causing SOD1 mutations can form similar amyloid fibril structures (Fig. 4) and perform similar functions implicated in ALS (Fig. 3). In sharp contrast, three genetic prion disease−related mutations E196K, F198S, and Y145Stop can form distinct amyloid fibril structures^46, 60–62^, highlighting the phenotypic diversity of these familial mutations in prion protein in an amyloid state.

Furukawa and co-workers described an alternative model of H46R fibrils and G85R fibrils based on protease digestion experiments and mass spectrometric analyses^23^. The so-called “core region model” predicts that the H46R fibril core contains one N-terminal segment comprising residues 1−30 and two C-terminal segments comprising residues ∼85−120 and ∼130−153, whereas the G85R fibril core contains one N-terminal segment comprising residues 1−30 and one C-terminal segment comprising residues ∼82−153 (ref. ^23^). This is in good agreement with our model, wherein β1-b4 and β5-b7 for the H46R fibril core would correspond to the second and third segments in the H46R fibril core region model, and β1-b8 for the G85R fibril core would correspond to the second segment in the G85R fibril core region model^23^. Interestingly, the Hart laboratory determined the crystal structure of ALS-causing SOD1 mutation H46R, and proposed that conformational change in H46R permits a gain of interaction between dimers that aggregate into zigzag filaments, highlighting the role of gain of interaction in pathogenic SOD1 (ref. ^63^). This is partly compatible with our model, wherein the H46R fibril core exhibits a serpentine fold containing seven β-strands.

Ferroptosis, a recently discovered iron-dependent form of regulated cell death^7, 9–11^, plays an important role in mediating selective motor neuron death in ALS^7–9^, but the mechanism behind the phenomenon remains poorly understood. The expressions of the master regulator of ferroptosis GPX4 are decreased in lumbar spinal cords of ALS mice and patients^7, 8^. Specifically, mitochondria in ferroptotic cells appear smaller than normal mitochondria with increased mitochondrial membrane density and reduction of mitochondria crista^10, 54, 55^. In this work, we report that GPX4 level in neuronal cells treated by H46R fibril seeds or G85R fibril seeds is significantly decreased compared with those treated by wild-type SOD1 fibril seeds. Morphologically, mitochondria undergoing ferroptosis show distinct changes compared to those in normal neuronal cells. We show that neuronal cells treated by H46R fibril seeds or G85R fibril seeds have about 30% more mitochondria undergoing ferroptosis than those treated by wild-type SOD1 fibril seeds. Overall, our results show that the fibril seeds from H46R and G85R significantly promote ferroptosis in neuronal cells in a similar way, compared to those from wild-type SOD1.

In summary, on the one hand, two ALS-causing SOD1 mutations, H46R and G85R, form similar novel amyloid fibril structures revealed by cryo-EM; the H46R fibril and the G85R fibril consist of a single protofilament with a fibril core comprising residues 85−153 or 82−153, respectively. On the other hand, the fibril seeds from H46R and G85R have a significantly higher ability to cause mitochondrial impairment and promote ferroptosis in neuronal cells compared to the wild-type fibril seeds. We find a direct link between amyloid fibrils formed by genetic mutations of SOD1 and GPX4-regulated ferroptosis implicated in ALS. The reported cryo-EM structure of the H46R/G85R fibril reveals an unusual overall structure when compared to the wild-type fibril, characterized by a disruption of crucial salt bridges, a C-terminal fibril core, an unidentified density flanking the protofilament in the G85R fibril, three hydrophilic cavities, and seven or eight instead of thirteen β strands in the core. The fibril structures will be valuable to understanding the structural basis underlying the degeneracy properties of familial mutations in an amyloid state and inspiring future research on the molecular mechanisms by which mutations in SOD1 promote ferroptosis and cause the familial form of ALS.

## Methods

### Protein purification

A plasmid-encoding, full-length human SOD1 (1−153) was a gift from Dr. Thomas O’Halloran (Chemistry of Life Processed Institute, Northwestern University). The sequence for SOD1 1−153 was expressed from the vector pET-3d, and two SOD1 mutants H46R and G85R were constructed by site-directed mutagenesis using a wild-type SOD1 template; the primers are shown in Extended Data Table 1. All SOD1 plasmids were transformed into *E. coli* BL21 (DE3) cells (Novagen, Merck, Darmstadt, Germany. Recombinant full-length wild-type human SOD1 and its variants H46R and G85R were expressed from the vector pET-30a (+) in *E. coli* BL21 (DE3) cells. SOD1 proteins wer purified to homogeneity by Q-Sepharose chromatography as describe by Chattopadhyay et al.^64^ and Xu et al.^65^. After purification, recombinant wild-type SOD1 and its variants H46R and G85R were demetallated by dialysis in 10 mM EDTA and 10 mM NaAc buffer (pH 3.8) five times as described by Chattopadhyay et al.^64^ and Xu et al.^65^. In all, 10 mM NaAc buffer (pH 3.8) and 20 mM tris-HCl buffer (pH 7.4) were used for further dialysis. The apo forms of wild-type SOD1, H46R, and G85R were then concentrated, filtered, and stored at −80 °C. AAnalyst-800 atomic absorption spectrometer (PerkinElmer) was used to quantify metal content of SOD1 samples. Samples of wild-type SOD1, H46R, and G85R contained <5% of residual metal ions, indicating that the samples were indeed in the apo state. SDS-PAGE and mass spectrometry were used to confirm that the purified apo SOD1 proteins were single species with an intact disulfide bond. A NanoDrop OneC Microvolume UV-Vis Spectrophotometer (Thermo Fisher Scientific) was used to determine the concentration of apo SOD1 proteins according to their absorbances at 214 nm with a standard calibration curve drawn by BSA.

### SOD1 fibril formation

The apo forms of recombinant full-length wild-type human SOD1 and its variants H46R and G85R were incubated in 20 mM tris-HCl buffer (pH 7.4) containing 5 mM TCEP and shaking at 37 °C for 40−48 h, and the SOD1 fibrils were collected. Large aggregates in SOD1 fibril samples were removed by centrifugation for 5000 × *g* at 4 °C for 10 min. The supernatants were then concentrated to ∼30 μM in a centrifugal filter (Millipore). A NanoDrop OneC Microvolume UV-Vis Spectrophotometer (Thermo Fisher Scientific) was used to determine the concentrations of the wild-type SOD1 fibril, the H46R fibril, and the G85R fibril according to their absorbances at 214 nm with a standard calibration curve drawn by BSA.

### TEM of H46R fibrils and G85R fibrils

H46R fibrils and G85R fibrils were examined by TEM of negatively stained samples. Ten microliters of SOD1 mutation fibril samples (∼30 μM) were loaded on copper grids for 30 s and washed with H_2_O for 10 s. Samples on grids were then stained with 2% (w/v) uranyl acetate for 30 s and dried in air at 25 °C. The stained samples were examined using a JEM-1400 Plus transmission electron microscope (JEOL) operating at 100 kV.

### AFM of H46R fibrils and G85R fibrils

H46R fibrils and G85R fibrils were produced as described above. Ten microliters of SOD1 mutation fibril samples (∼30 μM) were incubated on a freshly cleaved mica surface for 2 min, followed by rinsing three times with 10 μl of pure water to remove the unbound fibrils and drying at room temperature. The fibrils on the mica surface were probed in air by the Dimension Icon scanning probe microscope (Bruker) with ScanAsyst mode. The measurements were realized by using a SCANASYST-AIR probe (Bruker) with a spring constant of 0.4 N/m and a resonance frequency of 70 kHz. A fixed resolution (256 × 256 data points) of the AFM images was acquired with a scan rate at 1 Hz and analyzed by using NanoScope Analysis 2.0 software (Bruker).

### Cryo-EM of H46R fibrils and G85R fibrils

H46R fibrils and G85R fibrils were produced as described above. An aliquot of 3.5 μl of ∼30 μM SOD1 mutation fibril solution was applied to glow-discharged holey carbon grids (Quantifoil Cu R1.2/1.3, 300 mesh), blotted for 3.5 s, and plunge-frozen in liquid ethane using a Vitrobot Mark IV. The grids were examined using a Glacios transmission electron microscope, operated at 200 kV, and equipped with a field emission gun and a Ceta-D CMOS camera (Thermo Fisher Scientific). The cryo-EM micrographs were acquired on a Krios G4 transmission electron microscope operated at 300 kV (Thermo Fisher Scientific) and equipped with a Bio-Quantum K3 direct electron detector (Gatan). A total of 8,680 movies for H46R fibrils and 6,366 movies for G85R fibrils were collected in super-resolution mode at a nominal magnification of ×105,000 (physical pixel size, 0.84 Å) and a dose of 18.75 e^-^ Å^-2^ s^-1^ (see Table 1). An exposure time of 3.2 s was used, and the resulting videos were dose-fractionated into 40 frames. A defocus range of −1.2 to −2.0 μm was used.

### Helical reconstruction

All image-processing steps, which include manual picking, particle extraction, 2D and 3D classifications, 3D refinement, and post-processing, were performed by RELION-3.1 (ref. ^51^).

For the H46R fibril, 63,230 fibrils were picked manually from 8,680 micrographs, and 1024- and 686-pixel boxes were used to extract particles by 90% overlap scheme. Two-dimensional classification of 1024-box size particles was used to calculate the initial twist angle. In regard to helical rise, 4.8 Å was used as the initial value. Particles were extracted into 400-box sizes for further processing. After several iterations of 2D and 3D classifications, particles with the same morphology were picked out. Local searches of symmetry in 3D classification were used to determine the final twist angle and rise value. The 3D initial model was a cylinder which was built by RELION helix toolbox; 3D classification was performed several times to generate a proper reference map for 3D refinement. Three-dimensional refinement of the selected 3D classes with appropriate reference was performed to obtain final reconstruction. The final map of the H46R fibril was convergent with a rise of 4.88 Å and a twist angle of −1.079°. Postprocessing was preformed to sharpen the map with a *B* factor of −78.38 Å^2^. On the basis of the gold standard Fourier shell correlation (FSC) = 0.143 criteria, the overall resolution was reported as 3.11 Å. The statistics of cryo-EM data collection and refinement is shown in Table 1.

For the G85R fibril, 37,434 fibrils were picked manually from 6,366 micrographs, and 1024- and 686-pixel boxes were used to extract particles by 90% overlap scheme. Two-dimensional classification of 1024-box size particles was used to calculate the initial twist angle. In regard to helical rise, 4.8 Å was used as the initial value. Particles were extracted into 400-box sizes for further processing. After several iterations of 2D and 3D classifications, particles with the same morphology were picked out. Local searches of symmetry in 3D classification were used to determine the final twist angle and rise value. The 3D initial model was a cylinder which was built by RELION helix toolbox; and due to the similarity of the density maps between the H46R fibril and the G85R fibril, we next used the H46R fibril density map with an initial low-pass filter of 30 Å as a reference map; 3D classification was performed several times to generate a proper reference map for 3D refinement. Three-dimensional refinement of the selected 3D classes with appropriate reference was performed to obtain final reconstruction. The final map of the G85R fibril was convergent with a rise of 4.87 Å and a twist angle of −1.058°. Postprocessing was preformed to sharpen the map with a *B* factor of −82.56 Å^2^. On the basis of the gold standard Fourier shell correlation (FSC) = 0.143 criteria, the overall resolution was reported as 2.97 Å. The statistics of cryo-EM data collection and refinement is shown in Table 1.

### Atomic model building and refinement

Coot 0.8.9.2 (ref. ^66^) was used to build de novo and modify the atomic models of the H46R fibril and the G85R fibril. The models with three adjacent layers were generated for structure refinement. The models were refined using the real-space refinement program in PHENIX 1.15.2 (ref. ^67^). All density map-related figures were prepared in Chimera1.15. Ribbon representation of the structure of SOD1 fibril was prepared in PyMol 2.3.

### Cell culture

SH-SY5Y neuroblastoma cells (catalog number GDC0210) were obtained from China Center for Type Culture Collection (CCTCC, Wuhan, China) and cultured in minimum essential media and in Dulbecco’s modified Eagle’s medium (Gibco, Invitrogen), supplemented with 10% (v/v) fetal bovine serum (Gibco), 100 U/ml streptomycin, and 100 U/ml penicillin in 5% CO_2_ at 37 °C.

### Ultrathin section TEM

SH-SY5Y neuroblastoma cells were cultured in 6-well plates in the minimum essential medium for 1 day and then cultured with 0 μM SOD1 fibril seeds, 10 μM wild-type SOD1 fibril seeds, 10 μM H46R fibril seeds, and 10 μM G85R fibril seeds, respectively, for 3 days, and cells cultured with 20 mM Tris-HCl buffer (pH 7.4) containing 5 mM TCEP as a negative control. After prefixation with 3% paraformaldehyde and 1.5% glutaraldehyde in 1 × PBS (pH 7.4), the cells were harvested and postfixed in 1% osmium tetroxide for 1 h using an ice bath; the samples were then dehydrated in graded acetone and embedded in 812 resins. Ultrathin sections of the cells were prepared using a Leica Ultracut S Microtome and negatively stained using 2% uranyl acetate and lead citrate. The doubly stained ultrathin sections of cells were examined using a JEM-1400 Plus transmission electron microscope (JEOL) operating at 100 kV. The TEM images were analyzed by using Origin Pro software version 8.0724 (Origin Laboratory), and *p* values were determined using two-sided Student’s *t*-test. All experiments were further confirmed by biological repeats.

### Western blotting

For analysis by western blotting, SH-SY5Y cells were cultured in 6-well plates in the minimum essential medium for 1 day and then cultured with 0 μM SOD1 fibril seeds, 10 μM wild-type SOD1 fibril seeds, 10 μM H46R fibril seeds, and 10 μM G85R fibril seeds, respectively, for 3 days, and cells cultured with 20 mM Tris-HCl buffer (pH 7.4) containing 5 mM TCEP as a negative control. The cells were harvested and resuspended in lysis buffer (pH 7.6) containing 1% Triton X-100, 50 mM Tris, 150 mM NaCl, 1 mM phenylmethanesulfonyl fluoride, and protease inhibitors (Beyotime) on ice for half an hour. The cell lysates were centrifuged at 10,000 *g* for 10 min. The supernatants were boiled in SDS-PAGE loading buffer for 15 min, then subjected to 12.5% SDS-PAGE, and probed with the following specific antibodies: rabbit anti-GPX4 monoclonal antibody (Abcam ab125066, 1:2,000) and mouse anti-β-actin (Beyotime AA128, 1:1,000). The amount of loaded protein was normalized using a BCA Protein Quantification kit (Beyotime). For calculating the amounts of GPX4 and β-actin, ImageJ software (NIH, Bethesda, MD) was used to assess the densitometry of the corresponding protein bands. The normalized amount of GPX4 in SH-SY5Y cells was calculated as the ratio of the density of GPX4 band over the density of β-actin band in cell lysates and expressed as the mean ± S.D. (with error bars) of values obtained in three independent experiments.

### Statistical analysis

The data shown for each experiment were based on at least three technical replicates, as indicated in individual figure legends (Fig. 3 and Extended Data Fig. 1c,d). Data are presented as mean ± S.D., and *p* values were determined using a two-sided Student’s *t*-test in Fig. 3 and Extended Data Fig. 1c,d. All experiments were further confirmed by biological repeats.

### Reporting summary

Further information on research design and experimental design is available in the Nature Research Reporting Summary linked to this article.

## Data availability

The cryo-EM density maps of human H46R fibril and human G85R fibril have been deposited in the Electron Microscopy Data Bank (EMDB) under accession codes EMD-35460 and EMD-35459, respectively. The atomic coordinates for human H46R fibril and human G85R fibril generated in this study are deposited in the Protein Data Bank (PDB) under accession codes PDB 8IHV and PDB 8IHU, respectively. Previously published structure 1HL4 is available from PDB. Biological materials are available on request. The source data including the statistical source data for Fig. 3i,j,l and Extended Data Fig. 1c,d and the uncropped gel images for Fig. 3k are provided with this paper. Other data are available upon reasonable request.

## Acknowledgements

Y.L. and C.L. acknowledge funding from the National Natural Science Foundation of China (nos. 32271326, 32071212, 82188101, and 32171236). Y.L. also acknowledges financial support from the National Natural Science Foundation of China (no. 31770833) and the Key Project of Basic Research, Science and Technology R&D Fund of Shenzhen (no. JCYJ20200109144418639). C.L. was also supported by the Science and Technology Commission of Shanghai Municipality (nos. 20XD1425000, 2019SHZDZX02, and 22JC1410400) and the Shanghai Pilot Program for Basic Research – Chinese Academy of Sciences, Shanghai Branch (Grant No. CYJ-SHFY-2022-005). L.-Q.W. acknowledges financial support from the National Natural Science Foundation of China (no. 32201040), China Postdoctoral Science Foundation (nos. 2021TQ0252 and 2021M700103), and the Fundamental Research Funds for the Central Universities (no. 2042022kf1047). W.L. acknowledges financial support from the National Natural Science Foundation of China (no. 82271524). L.Z. acknowledges financial support from the Key Project of Basic Research, Science and Technology R&D Fund of Shenzhen (no. JCYJ20200109144418639). Cryo-EM data were collected at the Core Facility of Wuhan University, China. We thank T. V. O’Halloran (Northwestern University) for the gift of the human SOD1 plasmid; D. Li (Cryo-EM Unit, Core Facility of Wuhan University) and Y. Zeng (Cryo-EM Unit, Core Facility of Wuhan University) for technical assistance with cryo-EM; W. Zou (College of Life Sciences, Wuhan University) for technical assistance with the TEM of ultrathin sections of cells; and R. He (College of Pharmacy, Jinan University) and Y. Wang (Institute of Biophysics, Chinese Academy of Sciences) for helpful suggestions.

## Author contributions

L.Z., W.L., C.L. and Y.L. supervised the project. L.-Q.W., C.L. and Y.L. designed the experiments. L.-Q.W., M.-Y. Z., H.-Y.Y., X.H. and J.C. purified the human SOD1 proteins, wild-type SOD1 fibrils, H46R fibrils, and G85R fibrils. L.-Q.W. and M.-Y. Z. cultured the cells and performed western blotting analyses and TEM of ultrathin sections of cells. L.-Q.W., M.-Y. Z., H.-Y.Y., and Z. W. performed AFM experiments. L.-Q.W., Y.M., M.-Y. Z., H.-Y.Y., X.-N.L., K.Z., W.X., and D.L. collected, processed, and/or analyzed cryo-EM data. L.-Q.W., Y.M., C.L. and Y.L. wrote the manuscript. All authors proofread and approved the manuscript.

## Competing interests

The authors declare no competing interests.

---

**Extended Data Fig. 1.**
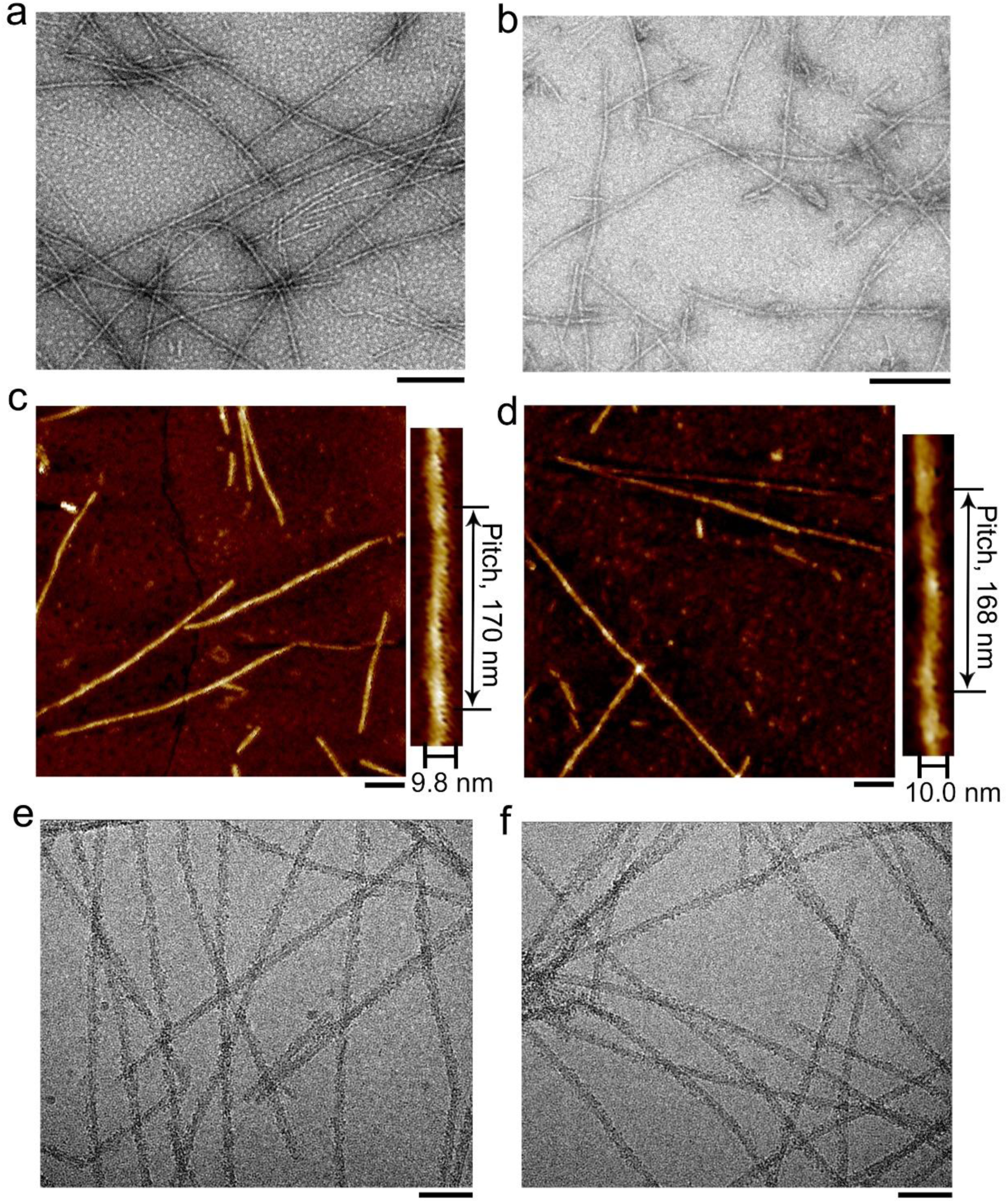
Comparison of the images of the H46R fibril and the G85R fibril. **a**,**b**, Negative-staining TEM images of amyloid fibrils produced from ALS-causing SOD1 mutations H46R (**a**) and G85R (**b**). **c**,**d**, AFM images of amyloid fibrils assembled from ALS-causing SOD1 mutations H46R (**c**) and G85R (**d**). Enlarged sections of **c** and **d** (right) showing the H46R fibril (**c**) and the G85R fibril (**d**) intertwined into a left-handed helix, with a fibril full width of 9.8 ± 1.0 nm and 10.0 ± 1.4 nm, respectively, and a helical pitch of 170 ± 9 nm and 168 ± 7 nm, respectively. The helical pitch and fibril width were measured and expressed as the mean ± SD of values obtained in *n* = 8 biologically independent measurements. Source data are provided as a Source Data file. **e**,**f**, Raw cryo-EM images of amyloid fibrils assembled from ALS-causing SOD1 mutations H46R (**e**) and G85R (**f**). The scale bars represent 200 nm (**a** and **b**) and 100 nm (**c** −**f**), respectively.

**Extended Data Fig. 2.**
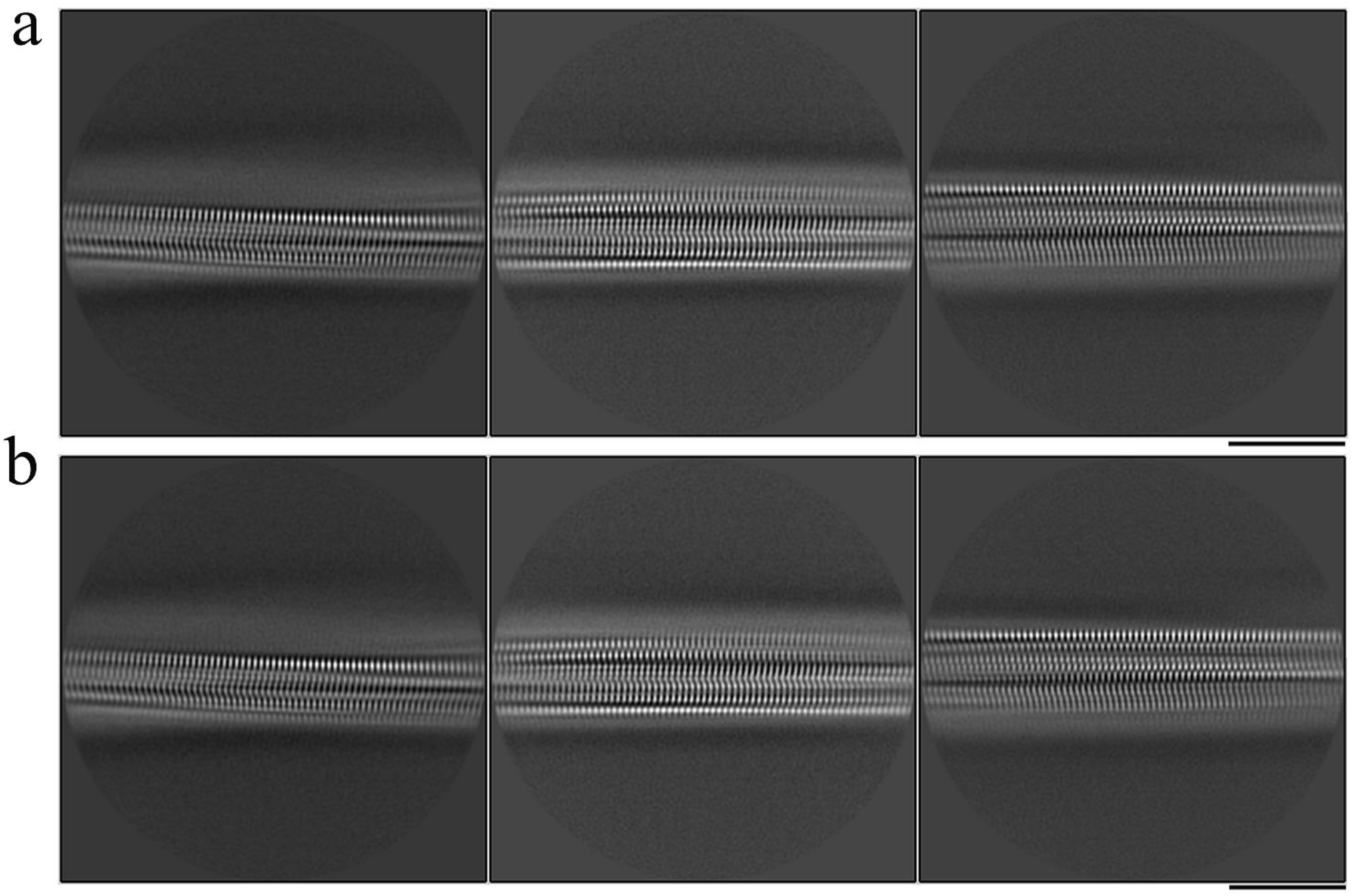
Comparison of the cryo-EM images of the H46R fibril and the G85R fibril. Reference-free 2D class averages of the H46R fibril (**a**) and the G85R fibril (**b**) both showing a single protofilament intertwined. Scale bar, 10 nm.

**Extended Data Fig. 3.**
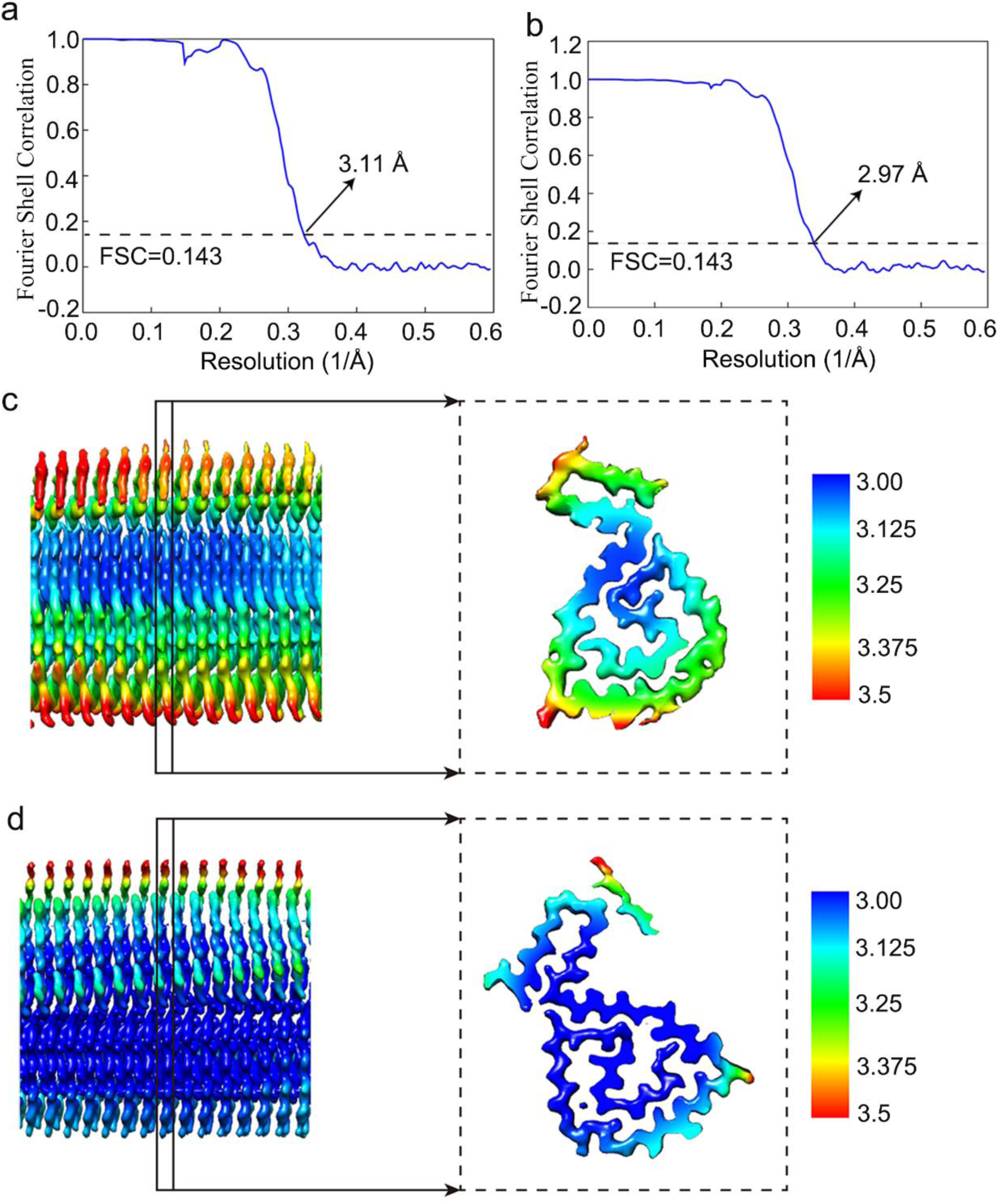
Global (a,b) and local resolution (c,d) estimates for the reconstructions of the H46R fibril and the G85R fibril. **a**,**b**, The reconstruction was reworked and gold-standard refinement was used for estimation of the density map resolution. The global resolutions of 3.11 Å for the H46R fibril (**a**) and 2.97 Å for the G85R fibril (**b**) were calculated using two Fourier shell correlation (FSC) curves (blue) cut-off at 0.143. **c**,**d**, The density maps of H46R fibrils (**c**) and G85R fibrils (**d**) are colored according to local resolution estimated by ResMap. The enlarged cross sections show the left top view of the density maps of a single protofilament in the H46R fibril (**c**) and a single protofilament in the G85R fibril (**d**). The color keys on the right show the local structural resolution in angstroms (Å) and the colored maps indicate the local resolution ranging from 3.0 to 3.5 Å.

**Extended Data Fig. 4.**
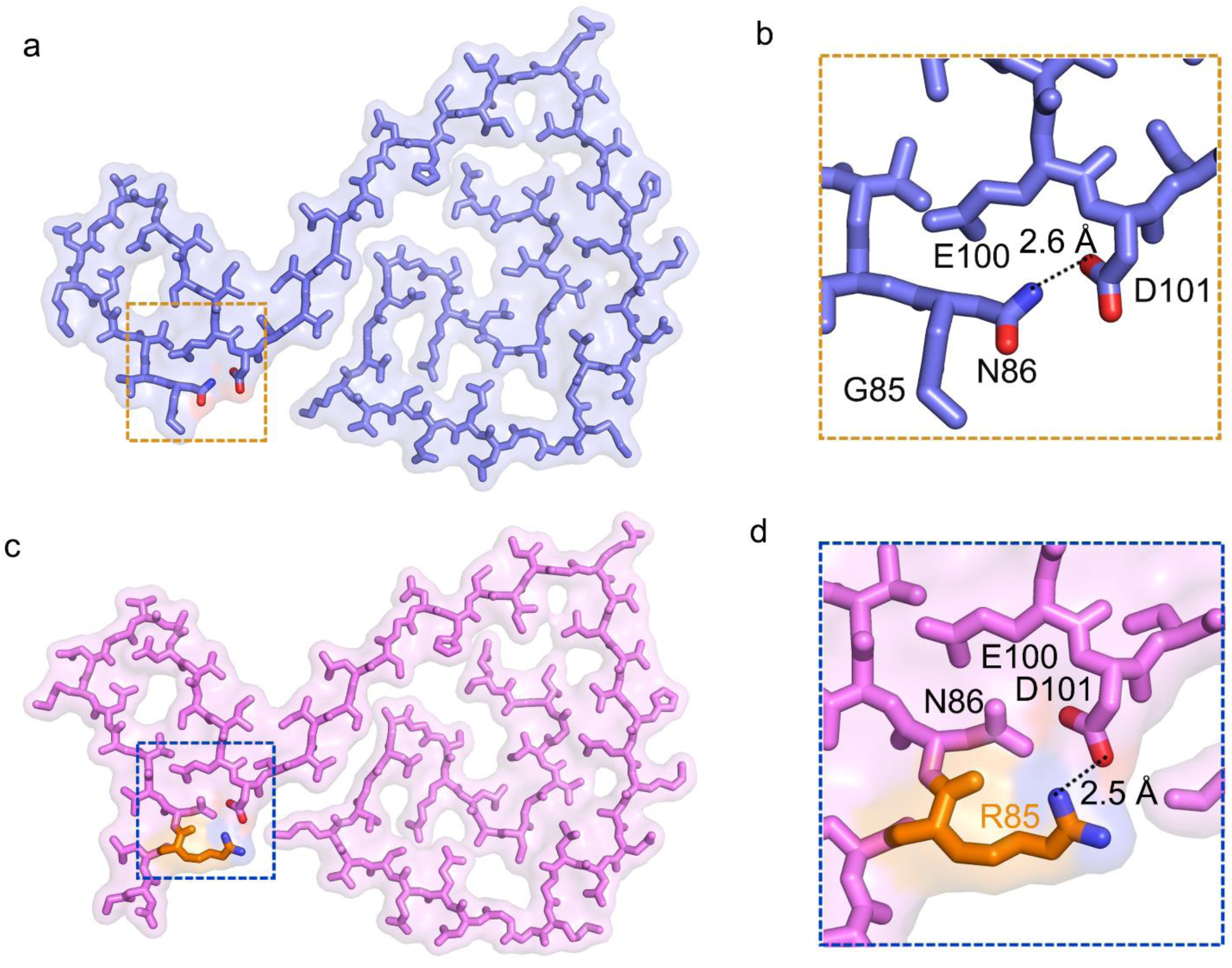
Close-up view of the stick representation of the structure of the H46R fibril stabilized by a hydrogen bond (a,b) and that of the G85R fibril stabilized by an intramolecular salt bridge (c,d). **a**, A space-filled model overlaid onto a stick representation of the H46R fibril in which a single protofilament is shown in blue. Asp/Asn pairs that form a hydrogen bond are highlighted in red (oxygen atom in Asp) and blue (nitrogen atom in Asn), and the hydrogen bond region is magnified in **b**. **b**, A magnified top view of the hydrogen bond region of an H46R protofilament, where a hydrogen bond is formed between Asp101 and Asn86, with a distance of 2.6 Å. **c**, A space-filled model overlaid onto a stick representation of the G85R fibril in which a single protofilament is shown in magenta. Asp/Arg pairs that form a new salt bridge are highlighted in red (oxygen atom in Asp) and blue (nitrogen atom in Arg), and the salt bridge region is magnified in **d**. **d**, A magnified top view of the salt bridge region of an G85R protofilament, where a strong salt bridge is formed between Asp101 and Arg85, with a distance of 2.5 Å. Arg85 in G85R variant is highlighted in orange.

**Extended Data Fig. 5.**
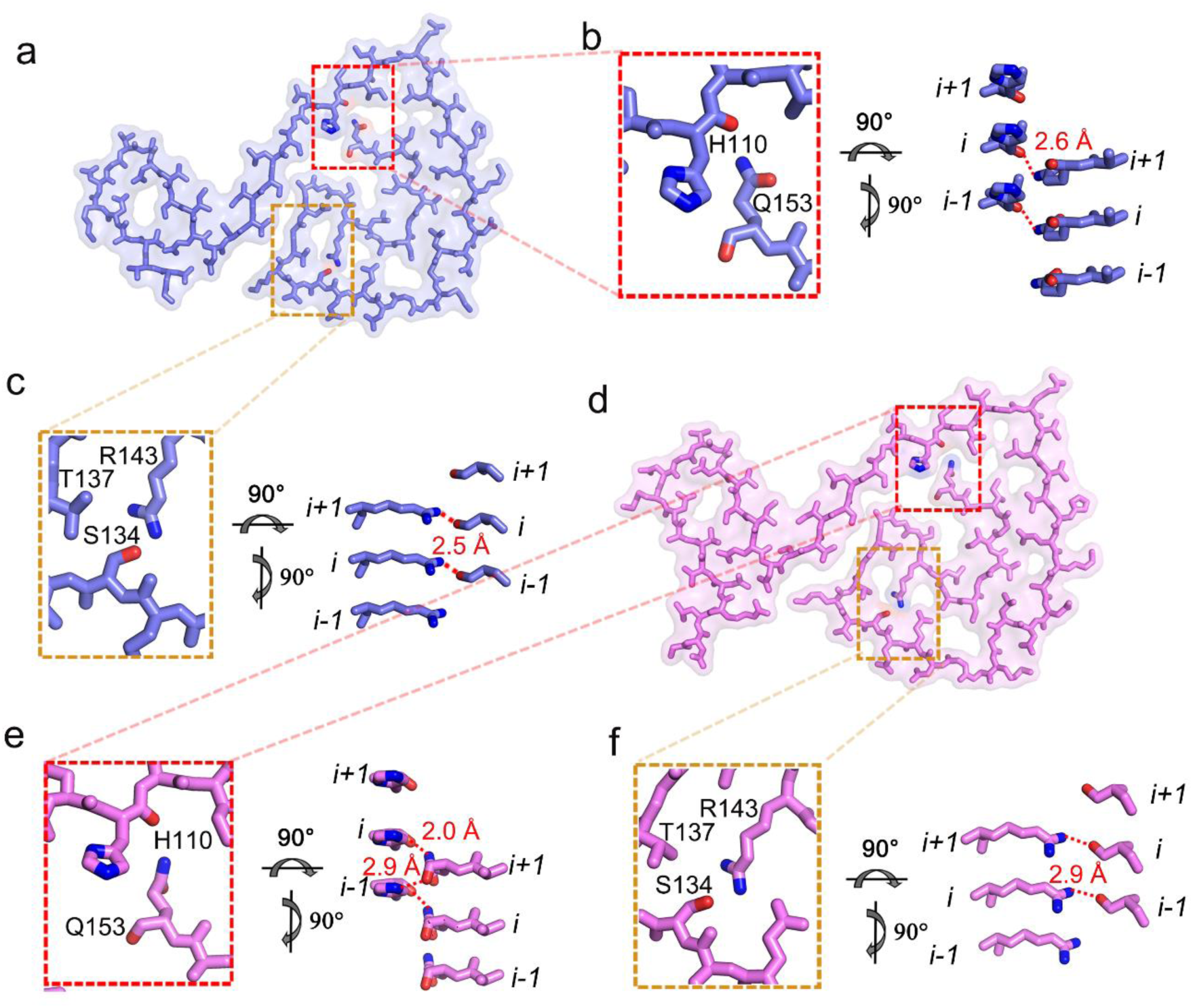
Close-up view of the stick representation of the structure of the H46R fibril stabilized by two hydrogen bonds (a−c) and that of the G85R fibril stabilized by three hydrogen bonds (d−f). **a**, A space-filled model overlaid onto a stick representation of the H46R fibril in which a single protofilament is shown in blue. His/Gln pairs and Ser/Arg pairs that form hydrogen bonds are highlighted in red (oxygen atoms in His and Ser) and blue (nitrogen atoms in Gln and Arg), and two hydrogen bond regions are magnified in **b**,**c**. **b**,**c**, Magnified top views of the two hydrogen bond regions of of an H46R protofilament, where two pairs of amino acids (His110 and Gln153; and Ser134 and Arg143) form two hydrogen bonds. Two side views (right) highlighting a hydrogen bond between the main chain of His110 from the molecular layer (*i*) and Gln153 from the adjacent molecular layer (*i* + 1), with a distance of 2.6 Å (red), or between Ser134 from the molecular layer (*i*) and Arg143 from the adjacent molecular layer (*i* + 1), with a distance of 2.5 Å (red). **d**, A space-filled model overlaid onto a stick representation of the G85R fibril in which a single protofilament is shown in magenta. His/Gln pairs and Ser/Arg pairs that form hydrogen bonds are highlighted in red (oxygen atoms in His and Ser) and blue (nitrogen atoms in Gln and Arg), and three hydrogen bond regions are magnified in **e**,**f**. **e**,**f**, Magnified top views of the three hydrogen bond regions of an G85R protofilament, where two pairs of amino acids (His110 and Gln153; and Ser134 and Arg143) form three hydrogen bonds. Two side views (right) highlighting a hydrogen bond between the main chain of His110 from the molecular layer (*i*) and Gln153 from the adjacent molecular layer (*i* + 1), with a distance of 2.0 Å (red), or between His110 from the molecular layer (*i* −1) and the main chain of Gln153 from the molecular layer (*i* + 1), with a distance of 2.9 Å (red), or between Ser134 from the molecular layer (*i*) and Arg143 from the adjacent molecular layer (*i* + 1), with a distance of 2.9 Å (red).

**Extended Data Table 1.**
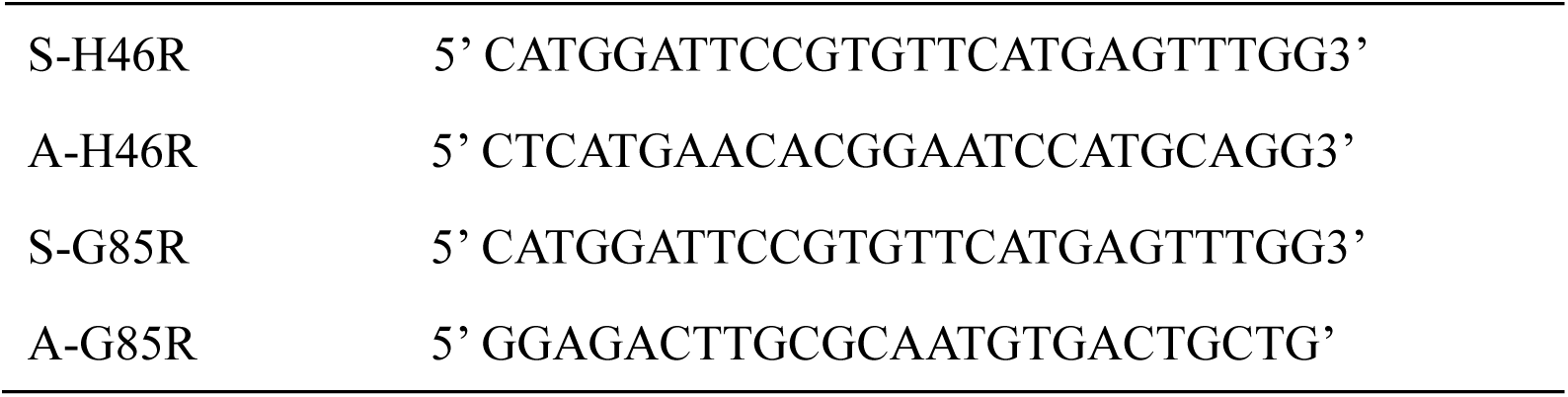
The primers designed for full-length human SOD1 with H46R mutation or G85R mutation

## Notes

### Competing Interest Statement

The authors have declared no competing interest.

